# Species-specific prophage induction by ciprofloxacin in human gut metagenomes

**DOI:** 10.64898/2026.03.11.711154

**Authors:** Ben Sakdinan, Anshul Sinha, Firdausi Qadri, Ashraful I. Khan, Eric J. Nelson, B. Jesse Shapiro

## Abstract

Antibiotics are known to trigger prophage induction in controlled laboratory settings, but it remains unclear whether this also occurs within microbiomes in nature. Current methods investigating the link between antibiotics and prophage induction within the human gut rely on *in vitro* culturing of human gut bacterial isolates. Using a metagenomic approach, we aimed to measure prophage induction and whether it is associated with antibiotic exposure. Across two independent human cohorts, we compared prophage to bacterial host read depth ratios (P:H) across known or measured antibiotic exposures. We found that induction is not broadly associated with antibiotic exposures at the level of the overall microbiome, but that ciprofloxacin increases P:H ratios in specific bacterial species. We documented heterogeneous trajectories of P:H ratios over the course of antibiotic exposure, sometimes increasing and remaining high, or returning to baseline. This study complements experimental models by providing *in vivo* evidence of induction in the human gut.

**Importance:** Bacteriophages are viruses that infect a bacterial host. The lytic and lysogenic cycles are the two classic outcomes of phage infection. In the lytic cycle, the phage immediately replicates and lyses its host to release new viral particles. In the lysogenic cycle, the phage, now called a prophage, integrates its genome into that of its host without killing it. Prophages can switch to the lytic cycle in a process called induction, in which the viral genome is replicated, the host cell is lysed, and viral particles are released. The most immediate consequence of induction is host cell death which can impact bacterial populations and communities. Since prophages are mobile genetic elements that can move between bacteria, they are also an important vehicle for horizontal gene transfer. While induction has been well studied *in vitro*, whether and how induction occurs within the complex microbial ecosystem in humans is less well characterized. Understanding prophage induction *in vivo* is therefore critical in corroborating *in vitro* observations.

## Main text

Current studies investigating the triggers of prophage induction employ *in vitro* or animal models,^1^ with little research directly studying the causes of induction within microbiomes. For example, ciprofloxacin triggers prophage induction *in vitro*, yet evidence of this occurring in humans is absent.^2–5^ Here we aim to address this knowledge gap by inferring prophage induction in metagenomic data from two human cohorts with varying levels of antibiotic exposure: diarrheal patients from Bangladesh with antibiotic exposures quantified in their stool by mass spectrometry, and healthy American (US) volunteers sampled before and after a course of ciprofloxacin. Using these datasets, we calculated the prophage to bacterial host read depth ratio (henceforth abbreviated to P:H ratio) as a signal for induction.

### No evidence for systematic community-wide prophage induction by antibiotics

To test the hypothesis that antibiotics systematically induce prophages in the gut microbiome, we mapped metagenomic reads to virus-containing contigs to measure the ratio of read depth of the prophage region to the host region. When a prophage is induced, its genome is replicated and released from the host cell as newly assembled virions. Such relative changes in genome copy number are captured by the P:H ratio (**Figure 1A**; Methods).^6^

**Figure 1.**
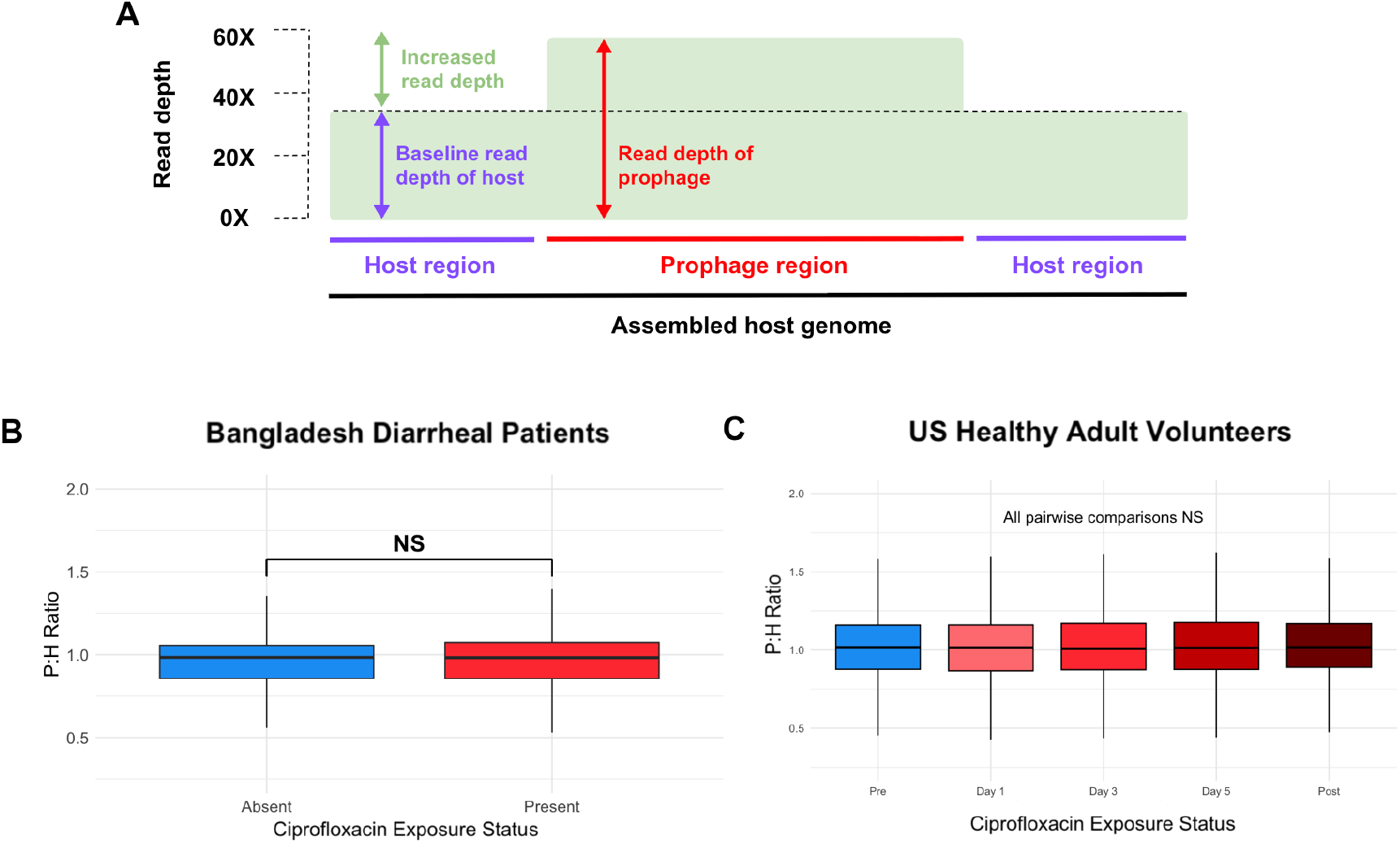
Quantification of prophage induction in human gut microbiomes across antibiotic exposures using metagenomic read mapping. (A) Conceptual visualization of the prophage to host (P:H) ratio as calculated from metagenomic reads mapped to assembled contigs containing prophages. (B) P:H ratio across ciprofloxacin exposure status in the Bangladesh cohort. Statistical significance was assessed with a Wilcoxon rank-sum test. (C) P:H ratios in healthy US volunteers before (pre), during (days 1, 3, and 5), and after (post) ciprofloxacin exposure. Statistical significance was assessed with a Wilcoxon rank-sum test; NS indicates Not Significant (*P* > 0.05).

We estimated the P:H ratio in 4055 prophages from the Bangladesh cohort and assessed their association with measured antibiotic exposures (**Supplementary Table 1**). Based on concentrations measured in our previous study, we set the following minimum concentrations to define biologically meaningful exposure to an antibiotic: ciprofloxacin (0.063 µg/mL), doxycycline (0.13 µg/mL), and azithromycin (8.0 µg/mL).^7^ We found no significant increase in community-wide P:H ratios in the presence of ciprofloxacin (**Figure 1B**) or other antibiotics. Azithromycin exposure was associated with a slight decrease in P:H ratios from a median of 1.0 to 0.98, potentially due to selection of bacterial genomes without an integrated prophage (Supplementary Material; **Figure S1**).^8^ As an additional test of our hypothesis in a healthy cohort, we calculated P:H ratios in 15,677 prophages assembled from United States healthy adult volunteers (Supplementary Table 1) and compared the ratios at timepoints before, during, and after a controlled ciprofloxacin exposure. We found no significant difference in P:H ratios from baseline (pre-exposure) to any post-antibiotic time point (**Figure 1C**). Together, the analysis of both cohorts indicates that antibiotic exposure is not measurably associated with broad-scale prophage induction in gut metagenomes.

### Ciprofloxacin is associated with prophage induction in specific bacterial species

While we found no evidence for measurable antibiotic-induced prophage induction at the overall community level, we hypothesized that induction could be restricted to specific taxa. To investigate this, we classified prophages by their predicted host taxonomy, and clustered them at the species or genus levels. To test this hypothesis in the Bangladesh cohort, we considered 19 prophages with an inferred bacterial host species that were present in at least 20 patients **(Figure S2**). Consistent with an experimental study,^4^ we found that ciprofloxacin exposure was associated with an increase in the median P:H ratio in *Salmonella enterica* from 0.94 to 1.13, and an increase in *Morganella morganii* from 1.02 to 1.09 (Wilcoxon rank-sum, *P* < 0.05 after a conservative Bonferroni multiple test correction). In both species, P:H ratios increased from ∼1 in the absence of ciprofloxacin exposure to >1 with detectable ciprofloxacin (**Figure 2A**). We detected no instances of significant decreases in P:H ratios with antibiotic exposure. More than one genetically distinct prophage was identified per host species in most patients, suggesting that these effects are driven by more than a single prophage per species (Supplementary Material; **Figure S3**). Induction is more likely driven by ciprofloxacin than by differences in patient disease status (Supplementary Material; **Figure S3**).

**Figure 2.**
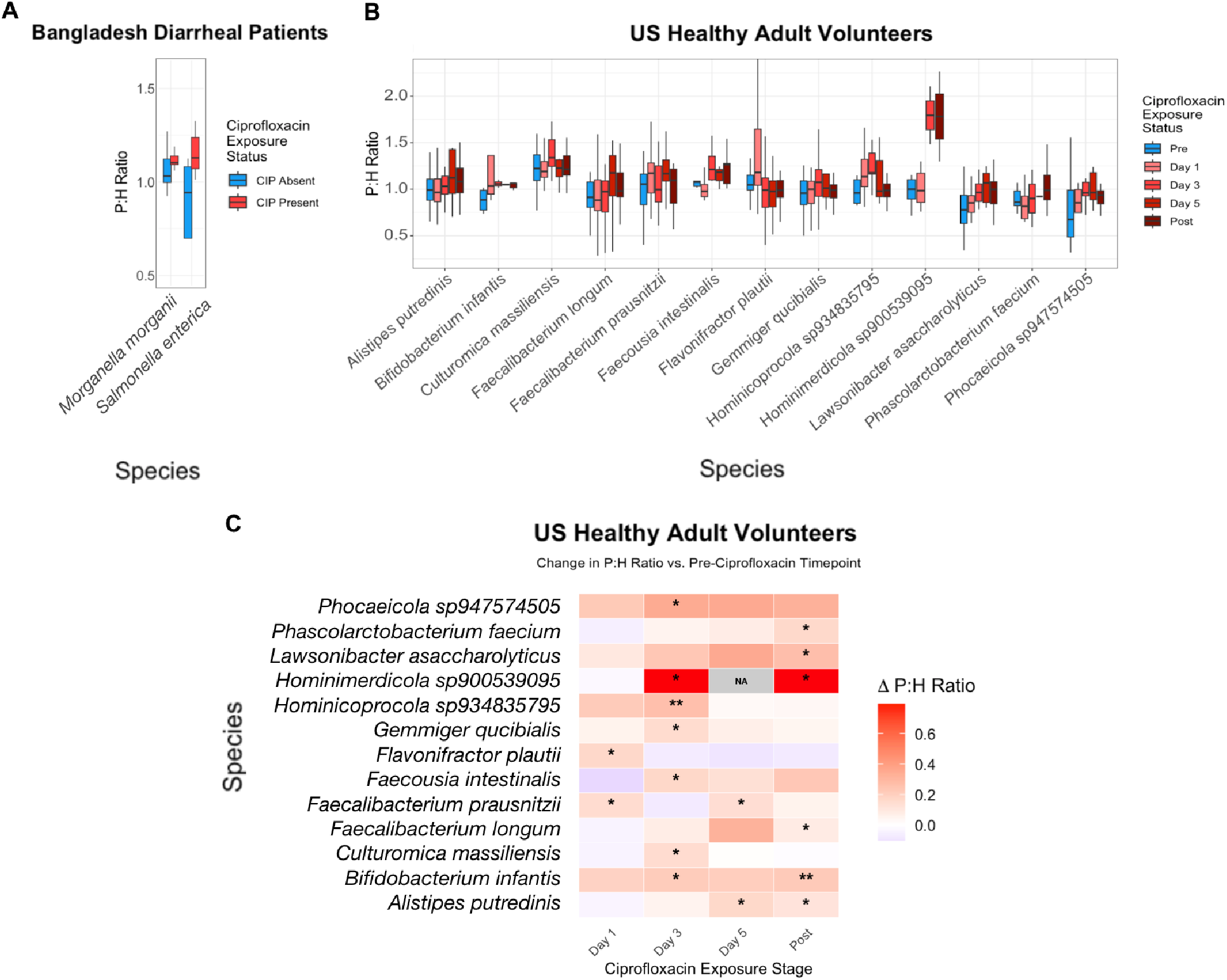
Ciprofloxacin exposure is significantly correlated with increased P:H ratio in multiple prophage host species. (A) Species in the Bangladesh cohort with a significant increase in P:H ratio between patients exposed vs. not exposed to ciprofloxacin. Statistical significance was assessed with a Wilcoxon rank-sum test with Bonferroni multiple correction. Only species with corrected *P* ≤ 0.05 are shown. (B) Species in the US healthy cohort with a significant increase in P:H ratios between baseline and a post-ciprofloxacin exposure time point. Only species with a Bonferroni corrected Wilcoxon rank-sum test *P*-value ≤ 0.05 at one or more time points are shown. (C) Heatmap showing the same species as in panel B. Cells are colored based on the difference (delta) between the P:H ratio at the given timepoint compared to baseline (pre-ciprofloxacin exposure). Bonferroni adjusted Wilcoxon rank-sum test *P-*values are coded as: NS: *P* > 0.05, *: *P* ≤ 0.05, **: *P* ≤ 0.01.

In the healthy US cohort, we considered 176 prophages matched with a host species and present in at least 20 patients. By comparing each post-exposure timepoint to their pre-exposure baseline, we found that ciprofloxacin exposure was significantly correlated with increased P:H ratios in at least one time point in 13 bacterial species (**Figure 2B**) and decreased P:H ratios in 5 species (Supplementary Material; **Figure S4**). Decreasing P:H ratios could be due to antibiotics selecting against lysogens relative to bacteria not containing prophages, reducing rates of induction, or both. We hypothesized that increased P:H ratios would lead to increased host bacterial lysis, reducing host relative abundance at subsequent time points. However, species with significantly increased P:H ratio at one post-exposure time point had no special tendency to decline in relative abundance at subsequent time points compared to the background distribution of species without evidence for induction (Kolmogorov-Smirnov test, *P* > 0.05), consistent with induction having only mild effects on host bacterial population sizes.^9,10^ The most common peak of induction was observed on the third day of ciprofloxacin exposure, with seven species having a significant difference at this timepoint (Wilcoxon rank-sum test, *P* < 0.05 after Bonferroni correction). Induction persisted at the post-exposure timepoint in six species (**Figure 2B-C**).

### Perspectives

We report evidence of species-specific prophage induction by ciprofloxacin in the human gut. The P:H ratios we measured are in line with other recent theory and metagenomic estimates,^9,10^ and lower than *in vitro* burst sizes yielding ratios of 100 or more.^11^ A variety of factors could yield this relatively low-level induction in the gut. Bacteria could experience lower antibiotic concentrations in the gut than typically applied in the lab. P:H ratios could also be suppressed when not all host genomes encode a prophage. An important caveat is that P:H ratios could be increased if antibiotics kill bacteria while free viral particles persist at stable abundances. Our interpretation of elevated P:H ratios as evidence for induction may therefore not be universal, but is a likely explanation given that ciprofloxacin is a known inducer of *Salmonella* and other bacteria under laboratory conditions. Regardless of the explanation, it is clear that antibiotics are not strong or universal inducers of prophages in the gut, but they likely have subtle, species-specific effects on P:H ratios. The generality of these results across antibiotic classes and host bacteria remain to be seen, along with potential consequences on microbiome structure and function.

## Methods

### Datasets

Publicly available sequencing data from our previous study was used to assemble 344 metagenomes from stool samples that were collected from cholera patients enrolled in hospitals across Bangladesh between March to April 2019.^7^ Dehydration severity for each patient was assessed and recorded as mild, moderate, or severe. This dataset contains concentrations of ciprofloxacin, doxycycline, and azithromycin, as measured by liquid chromatography mass spectrometry. Short reads for 344 cholera patients were accessed from our previous study by Madi et al. 2024,^7^ available in the NCBI Short Read Archive under BioProject PRJNA976726.

For our second dataset we analyzed publicly available metagnomes from Yaffe et al.^12^ This dataset contains metagenomic reads sequenced from stool samples collected from 60 healthy adults over 20 weeks surrounding a 5-day exposure event to ciprofloxacin. Participants took 500mg of ciprofloxacin twice daily during these 5 days. For each of the 60 patients in this longitudinal study, data from 5 timepoints were selected: 3 and 0 days before ciprofloxacin exposure (baseline), days 3, and 5 during ciprofloxacin exposure, and 5 days post exposure.

Reads were downloaded from NCBI under BioProject PRJNA974858.

### Metagenomic Data Analyses

Short reads for Bangladeshi diarrheal patient metagenomes were assembled using MEGAHIT v1.2.9.^13^ Long reads for healthy US adult volunteer metagenomes were assembled using metaFlye v2.9.6.^14^ Assembled contigs less than 1kb were filtered with SeqKit v2.5.1.^15^ Prophage contigs were predicted with geNomad v1.8.1.^16^ Quality and completeness of prophage contigs were assessed using CheckV1.0.3.^17^ Prophage to host read ratios were calculated using PropagAtE v1.1.0.^6^ Prophages were clustered into viral operational taxonomic units (vOTUs) based on shared gene content using VContTACT2 v0.11.3.^18^ Probable prophage hosts were predicted using iPHoP v1.3.3.^19^

### Host Relative Abundance Calculations

To calculate if host species with significant signals of prophage induction have larger changes in relative abundance than host species without significant signals or prophage induction, we calculated the change in prophage host’s relative abundance between each timepoint. This analysis was only completed within the US cohort, since timepoints did not exist in our Bangladesh cohort. Next, we subset the host species into those 13 that we detected significantly increased P:H ratios in, and all other hosts as a non-induced background signal. We used the Kolmogorov-Smirnov test to determine if the changes in relative abundances within our subset of significantly induced hosts significantly deviate from that of our non-induced hosts. However, no significant difference was observed at any timepoint.

### Statistical Analyses

All statistical analyses and generation of plots was conducted using R, version 2023.12.1+402 (2023.12.1+402).

To test for significant differences in P:H ratios between antibiotic exposure statuses, distributions of P:H ratios were compared using the Wilcoxon rank-sum test (Mann Whitney U test). In the case where multiple comparisons were made, such as in testing multiple host species, Bonferroni multiple test correction was performed. *P*-values were assigned with the following scheme: *P* > 0.05 = NS, *P* ≤ 0.05 = *, *P* ≤ 0.01 = **, *P* ≤ 0.001 = ***.

To test if host species had a special tendency to decline in relative abundance at the subsequent time points following a significant increase in P:H ratio, the difference in relative abundance between each timepoint was calculated for each of these species. Next, we subset these host species into those that had a significant signal of prophage induction, and those that did not. The distributions of relative abundance differences were compared between these two subsets using the Kolmogorov-Smirnov test.

## Code Availability

Example code used in this project is available on GitHub: https://github.com/bensakdinan/Discovering-phages-from-metagenomes.git

## Acknowledgments

We thank the patients and volunteers for participating in the studies and providing samples, as well as the clinical and laboratory teams who collected samples. We are grateful to S. Flora and colleagues at the Institute of Epidemiology, Disease Control and Research (IEDCR), Ministry of Health and Family Welfare, Government of Bangladesh, who collaborated on the original clinical studies in Bangladesh. We thank members of the Nelson and Shapiro labs for discussions that improved the manuscript.

## Supplementary Material

### Supplementary text

#### Prophage induction is primarily driven by ciprofloxacin, not disease-related stress

We recognize that the Bangladesh cohort suffered from diarrheal disease, potentially causing stress to the microbiome that could also modulate prophage induction. We previously found that antibiotic exposure is associated with milder dehydration severity in this cohort.^7^ The median P:H ratio of *Salmonella enterica* prophages was increased by 21% in patients with mild dehydration (**Figure S3**). It is unlikely that mild dehydration is causing induction, but rather that antibiotics are driving both induction and milder disease. Consistent with this hypothesis, among the 13 prophages found in patients with mild dehydration, all were exposed to ciprofloxacin, while only two of the 11 prophages in patients with non-mild dehydration were exposed to ciprofloxacin (Fisher’s exact test, OR > 5, *P* = 4.2e-5). These results suggest that ciprofloxacin exposure, not dehydration status, is the dominant factor contributing to prophage induction in diarrheal patients.

### Supplementary Figures

**Supplementary figure 1.**
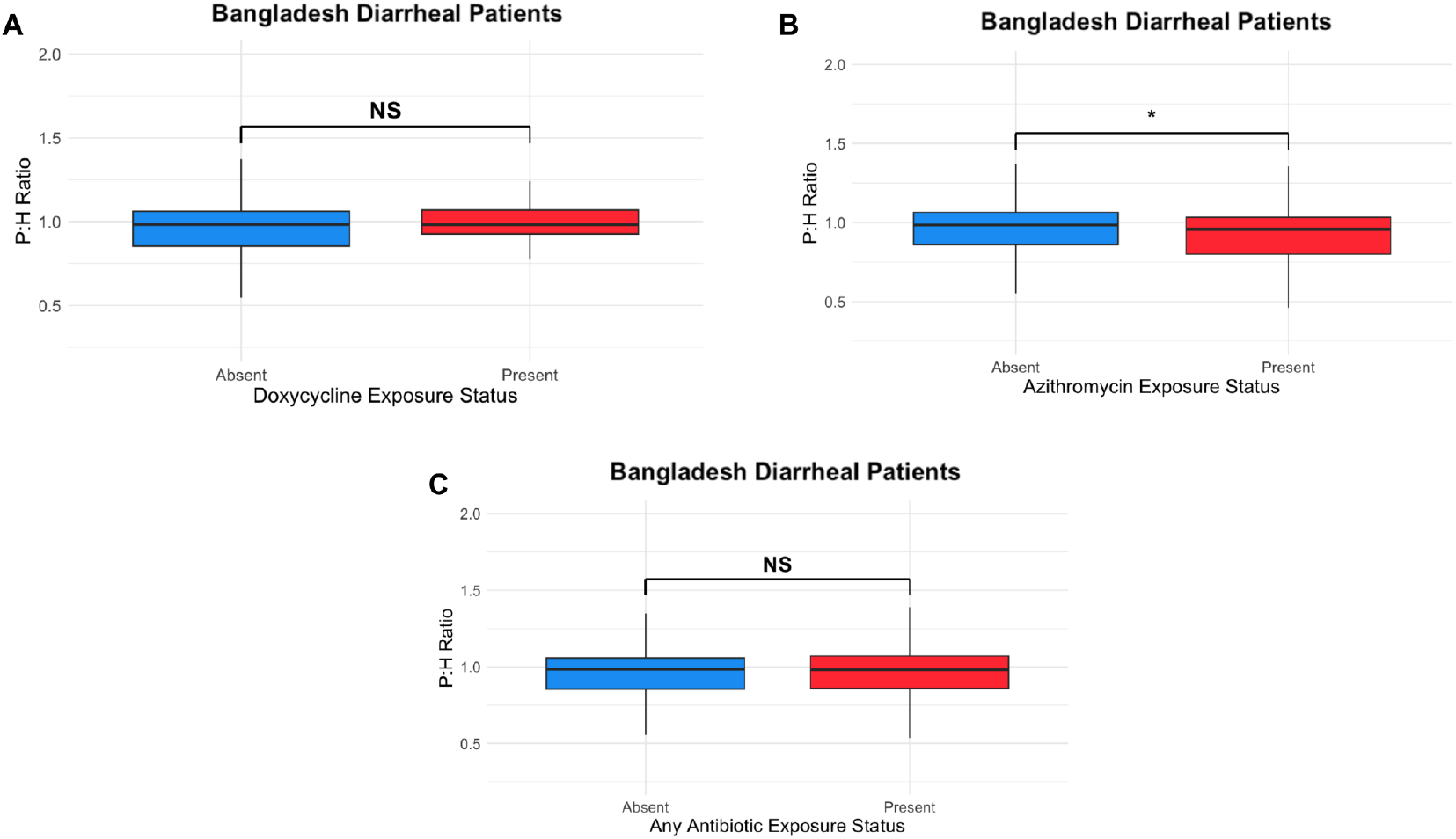
Signals of prophage induction across exposure to antibiotics and dehydration severity. (A) P:H ratio vs doxycycline exposure status within Bangladesh diarrheal patients. (B) P:H ratio vs azithromycin exposure status within Bangladesh diarrheal patients. (C) P:H ratio vs any antibiotic exposure status within Bangladesh diarrheal patients. Statistical significance was assessed with a Wilcoxon rank-sum test; NS: *P* > 0.05, *: *P* ≤ 0.05.

**Supplementary figure 2.**
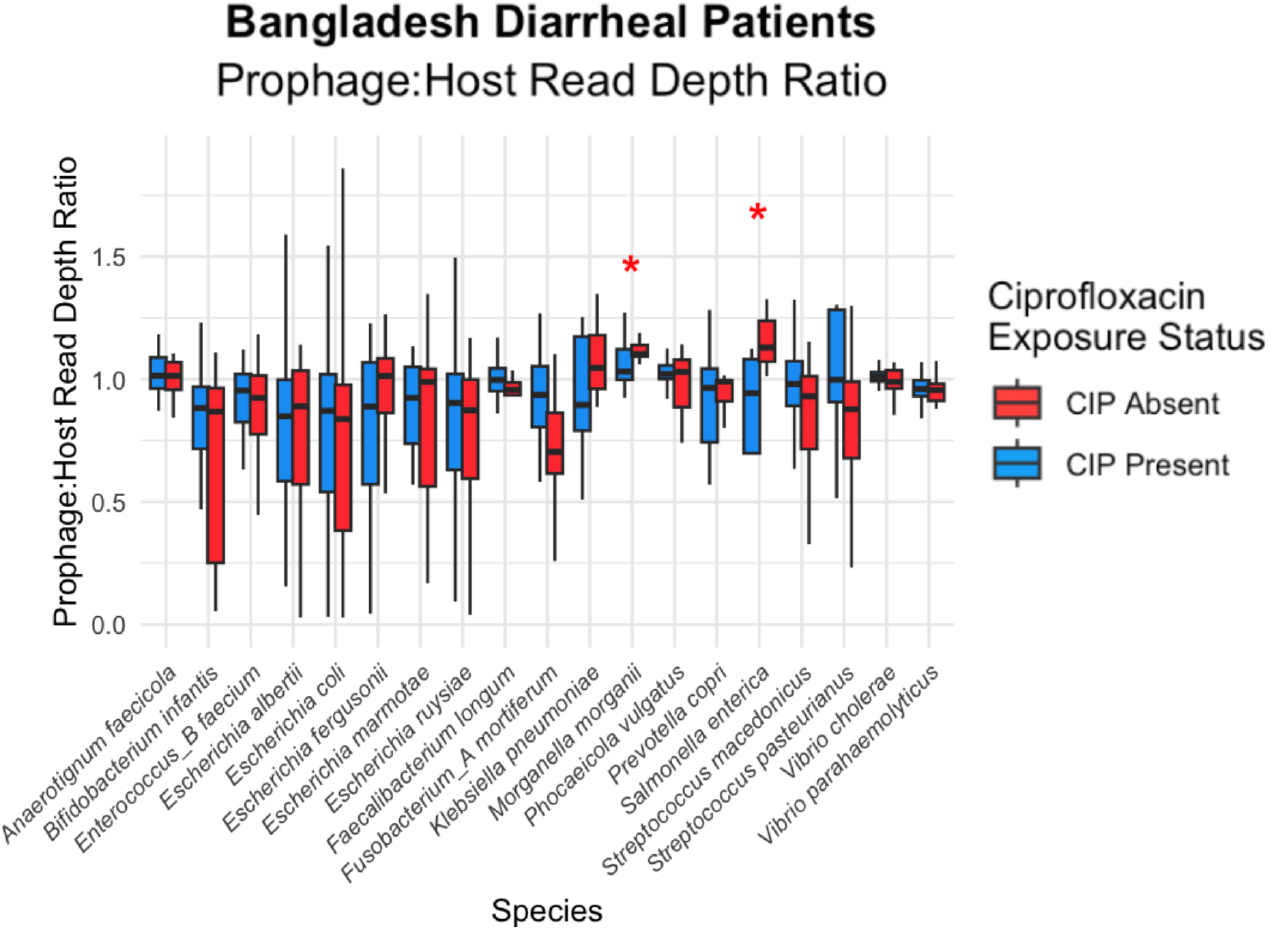
All species tested for induction in the Bangladesh dataset. P:H ratios vs ciprofloxacin exposure status in prophages clustered by their predicted host, within Bangladeshi diarrheal patients. To be included in this plot, a given species must have been assembled from at least 20 patients and contain a datapoint at both statuses of ciprofloxacin exposure. Statistical significance calculated by Wilcoxon rank-sum with Bonferroni multiple correction NS: *P* > 0.05, *: *P* ≤ 0.05.

**Supplementary figure 3.**
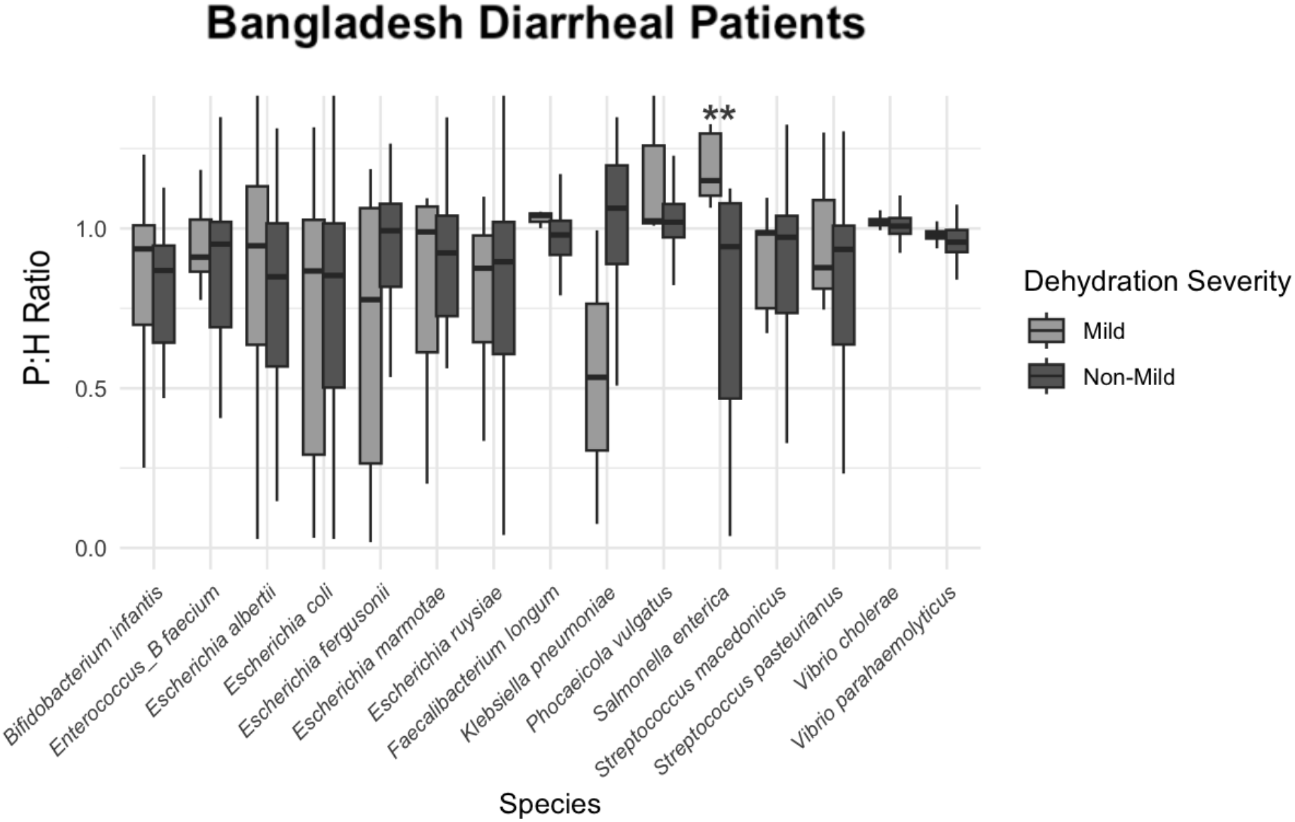
In Bangladeshi diarrheal patients, mild dehydration is significantly correlated with increased P:H ratio for *Salmonella enterica* prophages. P:H ratios vs dehydration severity in prophages clustered by their predicted host, within Bangladeshi diarrheal patients. Statistical significance calculated by Wilcoxon rank-sum with Bonferroni multiple correction NS: p > 0.05, *:*P* ≤ 0.05, **: *P* ≤ 0.01.

**Supplementary figure 4.**
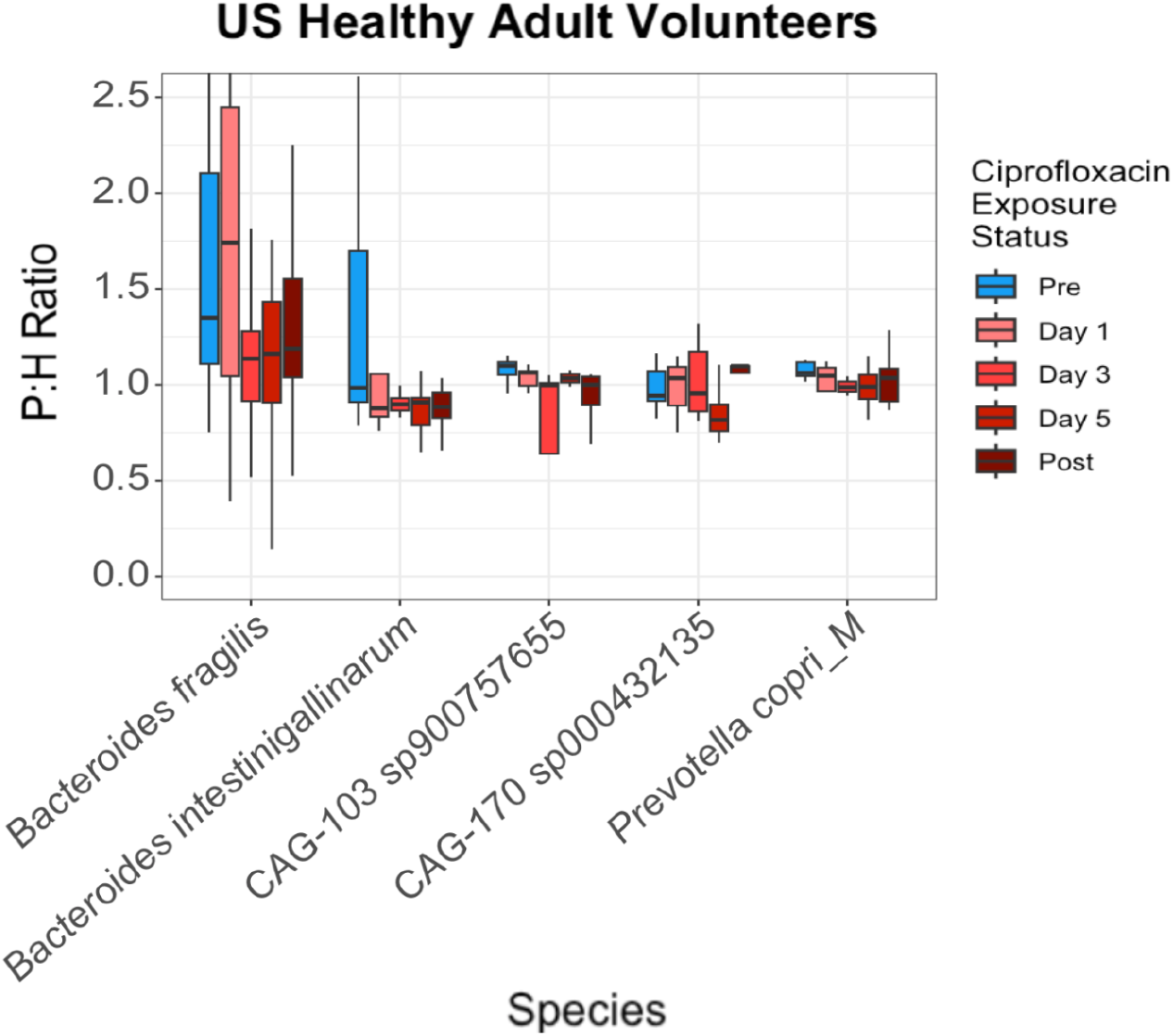
Ciprofloxacin exposure is significantly correlated with decreased P:H ratio in multiple prophage host species. Species in the US healthy cohort with a significant decrease in P:H ratios between baseline and a post-ciprofloxacin exposure time point. Only species with a Bonferroni corrected Wilcoxon rank-sum test *P*-value ≤ 0.05 at one or more time points are shown.

### Supplementary Table

**Supplementary Table 1.** Assembly of prophage contigs overview. Each row shows the number of contigs in each of the two datasets (columns) at each stage of assembly and annotation.

|  | *Bangladesh Diarrheal Patients* |  | *US Healthy Adult Volunteers* |  |
| --- | --- | --- | --- | --- |
|  | Total | Mean per sample | Total | Mean per sample |
| Contigs from metagenome assembly | 2,230,047 | 6,483 | 6,113,385 | 16,982 |
| Viral contigs | 233,184 | 678 | 686,520 | 1,907 |
| Prophage contigs | 4,055 | 12 | 15,677 | 44 |

## References

1. Balasubramanian, S., Osburne, M. S., BrinJones, H., Tai, A. K. & Leong, J. M. Prophage induction, but not production of phage particles, is required for lethal disease in a microbiome-replete murine model of enterohemorrhagic E. coli infection. PLoS Pathog. 15, e1007494 (2019).

2. Sutcliffe, S. G., Shamash, M., Hynes, A. P. & Maurice, C. F. Common Oral Medications Lead to Prophage Induction in Bacterial Isolates from the Human Gut. Viruses 13, (2021).

3. Goerke, C., Köller, J. & Wolz, C. Ciprofloxacin and trimethoprim cause phage induction and virulence modulation in Staphylococcus aureus. Antimicrob. Agents Chemother. 50, 171–177 (2006).

4. Braetz Sebastian, Schwerk Peter, Figueroa-Bossi Nara, Tedin Karsten, & Fulde Marcus. Prophage Gifsy-1 Induction in Salmonella enterica Serovar Typhimurium Reduces Persister Cell Formation after Ciprofloxacin Exposure. Microbiol. Spectr. 11, e01874–23 (2023).

5. Bucher, M. J., Puente, C. P., Sehdev, N. & Czyż, D. M. Sub-therapeutic Concentrations of Antibiotics Induce Prophage-driven Superinfection Exclusion and Fitness Cost in Pseudomonas aeruginosa. bioRxiv https://doi.org/10.1101/2024.11.20.624585 (2024) doi:10.1101/2024.11.20.624585.

6. Kieft Kristopher & Anantharaman Karthik. Deciphering Active Prophages from Metagenomes. mSystems 7, e00084–22 (2022).

7. Madi, N. et al. Phage predation, disease severity, and pathogen genetic diversity in cholera patients. Science 384, eadj3166.

8. Bailey, Z. M., Igler, C. & Wendling, C. C. Prophage maintenance is determined by environment-dependent selective sweeps rather than mutational availability. Curr. Biol. CB 34, 1739-1749.e7 (2024).

9. Lopez, J. A. et al. Abundance measurements reveal the balance between lysis and lysogeny in the human gut microbiome. Curr. Biol. CB 35, 2282-2294.e11 (2025).

10. Wirbel, J. et al. Long-read metagenomics reveals phage dynamics in the human gut microbiome. Nature https://doi.org/10.1038/s41586-025-09786-2 (2025) doi:10.1038/s41586-025-09786-2.

11. Abedon, S. T., Hyman, P. & Thomas, C. Experimental examination of bacteriophage latent-period evolution as a response to bacterial availability. Appl. Environ. Microbiol. 69, 7499–7506 (2003).

12. Yaffe, E. et al. Brief antibiotic use drives human gut bacteria towards low-cost resistance. Nature 641, 182–191 (2025).

13. Li, D., Liu, C.-M., Luo, R., Sadakane, K. & Lam, T.-W. MEGAHIT: an ultra-fast single-node solution for large and complex metagenomics assembly via succinct de Bruijn graph. Bioinforma. Oxf. Engl. 31, 1674–1676 (2015).

14. Kolmogorov, M. et al. metaFlye: scalable long-read metagenome assembly using repeat graphs. Nat. Methods 17, 1103–1110 (2020).

15. Shen, W., Sipos, B. & Zhao, L. SeqKit2: A Swiss army knife for sequence and alignment processing. iMeta 3, e191 (2024).

16. Camargo, A. P. et al. Identification of mobile genetic elements with geNomad. Nat. Biotechnol. https://doi.org/10.1038/s41587-023-01953-y (2023) doi:10.1038/s41587-023-01953-y.

17. Nayfach, S. et al. CheckV assesses the quality and completeness of metagenome-assembled viral genomes. Nat. Biotechnol. 39, 578–585 (2021).

18. Bin Jang, H. et al. Taxonomic assignment of uncultivated prokaryotic virus genomes is enabled by gene-sharing networks. Nat. Biotechnol. 37, 632–639 (2019).

19. Roux, S. et al. iPHoP: An integrated machine learning framework to maximize host prediction for metagenome-derived viruses of archaea and bacteria. PLoS Biol. 21, e3002083 (2023).

